# Characterisation of the faecal virome of captive and wild Tasmanian devils using virus-like particles metagenomics and meta-transcriptomics

**DOI:** 10.1101/443457

**Authors:** Rowena Chong, Mang Shi, Catherine E Grueber, Edward C Holmes, Carolyn Hogg, Katherine Belov, Vanessa R Barrs

**Affiliations:** School of Life and Environmental Sciences, University of Sydney, NSW 2006, Australia; Marie Bashir Institute for Infectious Diseases and Biosecurity, Sydney Medical School, University of Sydney, NSW 2006, Australia; School of Life and Environmental Sciences and Sydney Medical School, Charles Perkins Centre, University of Sydney, NSW 2006, Australia; San Diego Zoo Global, PO Box 120551, San Diego, CA 92112, USA; Sydney School of Veterinary Science, University of Sydney, NSW 2006, Australia

**Keywords:** Tasmanian devi, Marsupial, Carnivore, Gut microbiome, Endangered species, Virus

## Abstract

**Background:** The Tasmanian devil is an endangered carnivorous marsupial threatened by devil facial tumour disease (DFTD). While research on DFTD has been extensive, little is known about the viruses present in devils, and whether any of these are of potential conservation relevance for this endangered species.

**Methods:** Using both metagenomics based on virus-like particle (VLP) enrichment and sequence-independent amplification (VLP metagenomics), and meta-transcriptomics based on bulk RNA sequencing, we characterised and compared the faecal viromes of captive and wild Tasmanian devils.

**Results:** A total of 54 devil faecal samples collected from captive (n = 2) and wild (n = 4) populations were processed for virome characterisation using both approaches. We detected many novel, highly divergent viruses, including vertebrate viruses, bacteriophage and other dietary associated plant and insect viruses. In total, 18 new vertebrate viruses, including novel sapelovirus, astroviruses, bocaviruses, papillomaviruses and gammaherpesvirus were identified, as well as known mammalian pathogens including rabbit haemorrhagic disease virus 2 (RHDV2). Captive devils showed significantly lower levels of viral diversity than wild devils. Comparison of the two methodological approaches revealed substantial differences in the number and types of viruses detected, with meta-transcriptomics mainly identifying RNA viruses, and VLP metagenomics largely identifying DNA viruses.

**Conclusion:** This study has greatly expanded our knowledge of eukaryotic viruses in the Tasmanian devil and provides important baseline information that will contribute to the conservation and captive management of this endangered species. In addition, our results showed that a combination of VLP metagenomics and meta-transcriptomics may be a more comprehensive approach to virome characterisation than either method alone.

## Background

The Tasmanian devil (*Sarcophilus harrisii*) is the world’s largest extant carnivorous marsupial found only on the island state of Tasmania, Australia. Being predominantly a scavenger, the diet of devils largely comprises of carrion of mammals, such as wallabies, possums and kangaroos, although they are also known to consume and digest fish, insects, fruit and vegetation [1, 2].

Listed as endangered, the Tasmanian devil is facing the threat of extinction due to a contagious cancer, devil facial tumour disease (DFTD), that has caused drastic declines in wild devil populations by 77% since its discovery in 1996 [3]. There are currently two forms of DFTD affecting devils; DFT1 and DFT2 [4]. In an attempt to save the species from extinction, an insurance population was established in 2006 to serve as an important source of animals for supplementing wild populations at risk of population crashes due to DFTD [5, 6]. While extensive research has focused on DFTD itself as well as devil genetic diversity and susceptibility to the disease over the past decade [7-11], our understanding of other disease threats to devils remains limited. Specifically, virological studies are scarce and limited to the identification of a single gammaherpesvirus (DaHV-2), for which captivity was identified as a significant risk factor [12]. A comprehensive characterisation of the viral communities inhabiting the Tasmanian devil is an essential step to improving our understanding of host-microbe relationships, and maximising health and conservation management of the species.

Comparative analysis of marsupial viruses with those from diverse vertebrate hosts, including eutherian mammals, birds and other vertebrates will also provide a deeper understanding of the phylogenetic history of the viruses infecting this evolutionary unique group of mammals [13, 14].

To date, the most widely used method for studying viral metagenomics relies on the enrichment of virus-like particles (VLP) and subsequent sequence-independent amplification prior to sequencing [15-17]. Removal of non-viral genomic host and bacterial nucleic acids and enrichment of VLP is often necessary for the detection of low-titre viruses [17, 18]. More recently, the use of RNA sequencing of total non-ribosomal RNA from environmental samples gave rise to viral meta-transcriptomics, which has been successfully applied to characterise the viromes of diverse invertebrate and vertebrate species [13, 19, 20]. To our knowledge, no studies have been conducted to compare these two approaches, although doing so would allow us to understand the detection capabilities and biases associated with these different nucleic acid extraction and sample treatment methods.

We characterised the faecal virome of wild and captive Tasmanian devil using both the metagenomics approach based on VLP enrichment and sequence-independent amplification (hereafter “VLP metagenomics”) and the meta-transcriptomics approach based on RNA sequencing of total non-ribosomal RNA (hereafter “meta-transcriptomics”). Our objectives were threefold: (1) to provide a comprehensive characterisation of the faecal virome of Tasmanian devils; (2) to compare the two virome characterisation approaches, highlighting their advantages and potential challenges; and (3) to compare the faecal viromes of wild and captive Tasmanian devils.

## Materials and Methods

### Sample collection

Faecal samples were collected from wild Tasmanian devils between September 2016 and June 2017 from four locations in Tasmania (Figure 1); **A** Stony head (SH), **B** Buckby’s Road (BR), **C** Maria Island (MI), and **D** wukalina/Mt William National Park (wMW), as well as from captive devils at two Australian mainland zoos in June and July 2017 (Zoo A and Zoo B). Wild devils were trapped overnight during routine monitoring trips by Save the Tasmanian Devil Program staff, using baited PVC-pipe traps [21]. Upon capture, each animal was subjected to a thorough health check, including body weight measurement, estimation of body condition score and observation for signs of disease. Fresh faecal samples were collected from either the traps, or from the hessian bag, during processing of the devils. For captive devils, faeces were collected from the enclosures shortly after defecation. All samples were stored in either liquid nitrogen or a portable -80^°^C freezer (Stirling Ultracold, Global Cooling Inc.) immediately after collection. After arriving at the laboratory, samples were separated into two aliquots to be used in the extraction of total RNA for meta-transcriptomics and the enrichment of VLP for VLP metagenomics.

**Figure 1.**
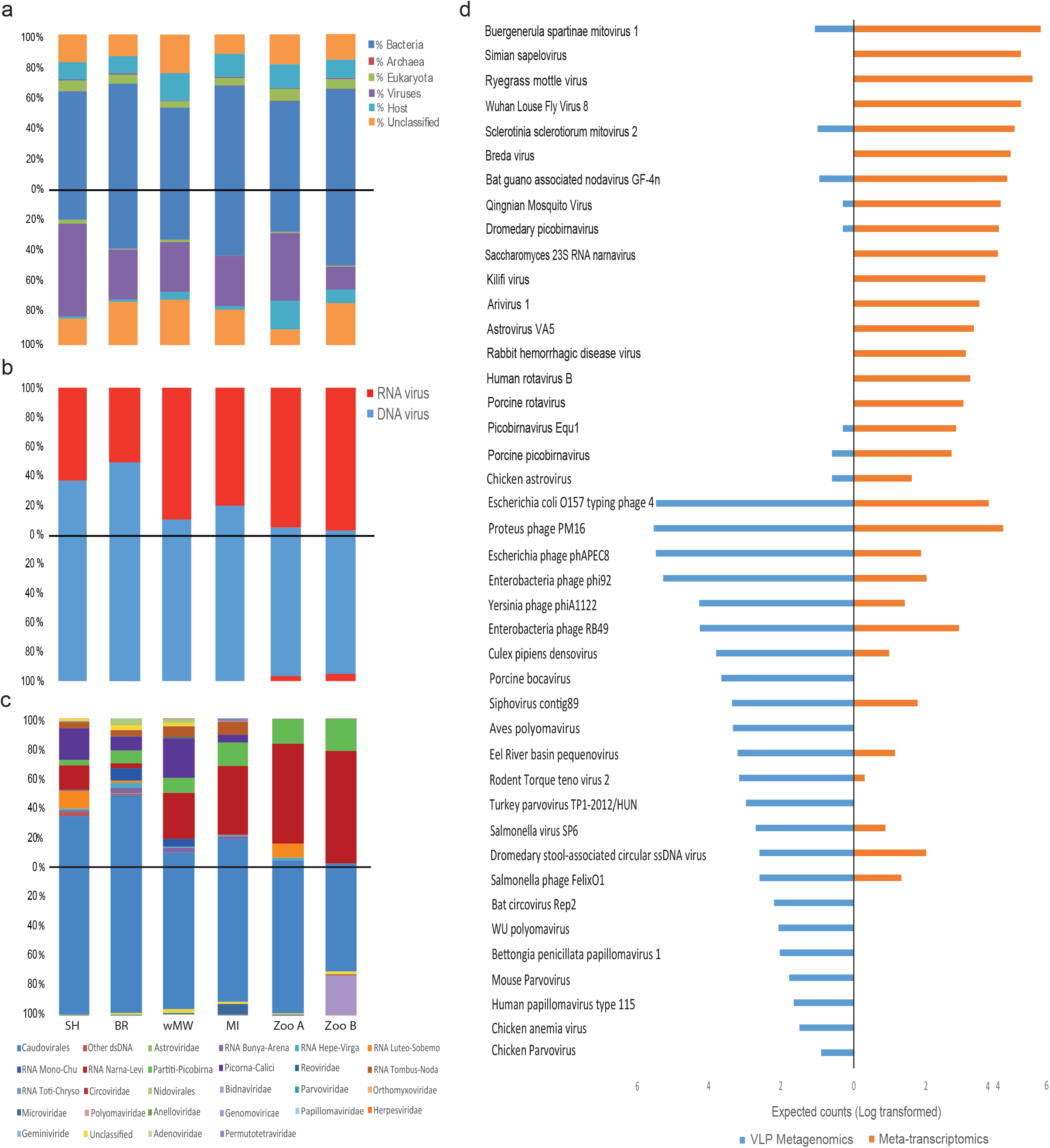
Map of Tasmania, Australia showing the four wild sampling sites.

### Meta-transcriptomics

#### Total RNA extraction, library preparation and sequencing

Samples were disrupted and homogenized in 600 μl of lysis buffer with 1.44 mm ceramic beads using a Bead Ruptor Homogenizer (Omni International) at 5 m.s^-1^ for 5 min. Total RNA was isolated using the Qiagen RNeasy Plus Mini Kit (Qiagen) following the manufacturer’s instructions. The extracted total RNAs were pooled based on their source locations at equal mass amount, with each pool containing 5-10 samples. The RNA pools were depleted of host and bacteria rRNA using a Ribo-Zero-Gold (Epidemiology) kit (Illumina) before constructing sequencing libraries using a TruSeq total RNA Library Preparation Kit (Illumina). Paired-end (75 bp) sequencing of each library was performed on a NextSeq500 HO platform (Illumina) at the Ramaciotti Centre for Genomics (Sydney, Australia).

#### VLP Metagenomics

##### VLP enrichment and nucleic acid extraction

A second aliquot from each faecal sample was processed for the VLP metagenomics approach, as described previously with minor modifications [17].

VLPs were enriched as follows: Faecal suspensions consisting of 50 mg faeces in 700 μl of were homogenized for 1 min using the Bead Ruptor Homogenizer (Omni International) at 5 m.s^- 1^ and centrifuged at 15000 × g for 3 min at 4 ^°^C. The resulting supernatants were filtered through 0.45 μm membrane filters (Corning) for 1 min at 15000 × g. The filtrates were treated with 0.8 μl benzonase (Sigma Aldrich), 7 μl Turbo DNase (Ambion), 1 μl micrococcal nuclease (New England Biolab) and 14.88 μl 1 x Turbo DNase buffer (Ambion) at 37 °C for 2 h to digest free-floating nucleic acids. Viral DNA and RNA were then simultaneously extracted using the QIAamp Viral RNA Mini Kit (Qiagen) according to the manufacturer’s instructions, except without addition of carrier RNA to the lysis buffer [17] .

##### Random amplification

Extracted viral nucleic acids were pooled based on their source locations at equal mass amounts, using the same representative samples as the total RNA preparations for meta-transcriptomics described above. The pooled extractions were subjected to first and second strand synthesis and random PCR amplification for 22 cycles using the Complete Whole Transcriptome Amplification (WTA2) kit (Sigma-Aldrich). The denaturation temperature was increased from 70°C to 95°C to allow for the denaturation and amplification of both dsDNA and RNA [17]. WTA2 PCR products were then purified using Agencourt AMPure XP beads (Beckman Coulter) prior to library preparation and sequencing.

##### Library preparation and Illumina Sequencing

Sequencing libraries were constructed using the Nextera XT DNA Library Preparation Kit (Illumina) according to the manufacturer’s instructions with the following modifications as shown in [17] : (i) the tagmentation time was reduced to 4 min to increase fragment size, (ii) input DNA was increased to 1.2 ng/ μl^-1^ and (iii) reagent quantities were halved to further increase the fragment size. Paired-end (100 bp) sequencing of each library was performed on a Hiseq2500 platform (Illumina) at the Ramaciotti Centre for Genomics (Sydney).

#### Assembly and annotation

Sequencing reads were de-multiplexed and quality trimmed with Trimmomatic [22] and assembled *de novo* using Trinity [23]. Resulting contigs were compared against the non-redundant nucleotide (nt) and protein (nr) databases on GenBank using Blastn and Blastx, respectively, with an e-value cut-off at 1E-5. Blast search using Blastx was also conducted against a bespoke database containing all viral RNA-dependent RNA polymerase (RdRp) protein reference sequences downloaded from GenBank. Taxonomic information at the domain level (i.e. Eukarya, Bacteria, or Archaea), as well as for viruses, was first assigned based on the blastn results and then based on blastx results. Potential virus-related sequences were further categorized into families and orders based on their genetic similarity to their closest relatives and/or their phylogenetic position. Similarly, the assignment of viruses to the broader groups of their hosts (i.e. Eukarya, Bacteria, or Archaea) was based on their phylogenetic relationship to viruses with reliable host information obtained either using experimental or phylogenetic approaches. The genetic identity cut-off for host assignment was set to be 40% based on the most conserved proteins such as RdRp or DNA polymerases [24]. The threshold was set based on the intra-family diversity of most vertebrate-specific virus families/genera [25]. The assignment of vertebrate host was based on phylogenetic analyses, in which a potential devil associated virus is expected to either cluster within, or form a sister group to, an existing mammalian virus group.

To compare the abundance of each transcript/contig, we calculated the percentage of total reads in the library. The abundance of host transcripts/contigs was estimated by mapping reads against Tasmanian devil genome using Bowtie2 [26], whereas those of other organisms, namely, viruses, bacteria, archaea, and non-host eukaryotes, were estimated using the RSEM approach [27] implemented in Trinity.

For each virus, the genome sequence was further extended by merging related contigs. Gaps in the genome were filled either by mapping the reads against the existing scaffold or by RT-PCR and Sanger sequencing. Putative ORFs in viral genomes were predicted by the Geneious 8.1 software [28] or NCBI ORF finder, and annotated based on their similarity to previous published virus genomes.

#### Phylogenetic analysis

Nucleotide sequences of complete or partial genomes and amino acid sequences from the conserved domain (e.g. the RNA-dependent RNA polymerase; RdRp) of the newly characterised viral sequences were aligned with those of reference viruses representative of the diversity of the corresponding virus family or species. The alignment was performed using the E-INS-I algorithm implemented in MAFFT (version 7) [29]. The quality of the alignments was subsequently assessed, and all ambiguously aligned regions were removed using TrimAl (version 1.2) [30]. Phylogenetic trees of aligned amino acid (all data sets with the exception of RHDV) or nucleotide (RHDV) sequences were then inferred using the maximum likelihood method implemented in PhyML version (3.0) [31], utilising the best-fit substitution model and the Subtree Pruning and Regrafting (SPR) branch-swapping algorithm.

#### Analyses of virome ecology

Viral abundance tables (Additional file 2: Table S2 and Additional file 3: Table S3) were generated based on complete or near complete viral contigs and the percentage of reads mapped to them using Bowtie 2 [26] in each library. QIIME (v 1.9) [32] was used to perform ecological and statistical analysis to compare viromes of different populations. Within-library virotype richness (alpha diversity measured using the *Chao1* metric) and dissimilarity between libraries (beta diversity measured using the Euclidean metric) for both VLP metagenomics and meta-transcriptomics were calculated based on levels of viral abundance. The statistical significance of differences in alpha diversity was evaluated by the Monte Carlo method (999 permutations), with a null hypothesis that diversity is equal in all libraries with a significance threshold of α = 0.05. Levels of viral abundance were also used to produce heatmaps and dendrograms from hierarchical clustering. Principal coordinates analysis (PCoA) was performed on the Euclidean distance matrix as calculated in QIIME, and additional cluster analysis was conducted using K-Means clustering in R [33].

#### PCR confirmation and Sanger sequencing of rabbit haemorrhagic disease virus (RHDV)

Contigs with high percentage of similarity (>97%) to rabbit haemorrhagic disease virus (RHDV) were detected in one of the meta-transcriptomics libraries. RHDV is used as a biocontrol for rabbits in Australia and causes fatal hepatitis in European rabbits (*Oryctolagus cuniculus*) and some hare species [34]. To confirm the detection of RHDV, RT-PCR was performed on individual faecal RNA extractions using the Qiagen OneStep Ahead RT-PCR kit (Qiagen) and primers from a previously validated Australian RHDV strain-specific PCR [35] as well as two additional primer sets manually designed based on the current meta-transcriptomics assembled contigs (Additional file 4: Table S4). PCR products were separated on 1.5% agarose gel (Bio-Rad Laboratories, Hercules, CA, USA) in 1x tris-acetate EDTA, and visualized using SYBR Safe DNA gel stain (Life Technologies, Carlsbad, CA, USA). Sanger sequencing of positive PCR products was performed at the Australian Genome Research Facility (Sydney, Australia). In addition, DNA was extracted from faecal samples using the ISOLATE Fecal DNA kit (Bioline, London, UK) and presence of rabbit DNA was tested using primers targeting a 110 bp region of the *Oryctolagus cuniculus* 12S mitochondrial rRNA gene (Fwd: 5′□CAAAAGTAAGCTCAATTACCACCGTAc3′; Rev: n5′□ATAAGGGCTTTCGTATATTCGGAAc3′) [36]. Rabbit DNA extracted from rabbit liver obtained from collaborators at CSIRO using the Bioline Isolate II Genomic DNA kit (Bioline) was used as a positive control. Cleanup, primer trimming and sequence analysis of Sanger data was performed using DNASTAR Lasergene [37] and Geneious [28].

## Results

### Overview of the devil virome

Location Captive/Wild Meta-transcriptomics No. reads No. contigs VLP metagenomics No.reads No. contigs We characterised the faecal virome of six pools consisting of a total of 54 faecal samples collected from Tasmanian devils from four wild and two captive sites using both meta-transcriptomics and VLP metagenomics approaches. Meta-transcriptomic libraries resulted in 128 - 140 million reads per pool (793,038,436 reads in total), which were assembled *de novo* into 196,919 – 358,327 contigs (Table 1). Blast analyses of sequence reads from the meta-transcriptomic protocol revealed large proportions of reads from Bacteria (55.11 – 67.77%), and only 4.32 – 7.88% from Eukarya. Mapping reads to the Tasmanian devil genome revealed that 10.75 – 18.99 % of reads originated from the host. The percentage of reads related to Archaea was less than 0.02%, and for viruses between 0.68 – 1.16% (Figure 2a).

**Table 1.**
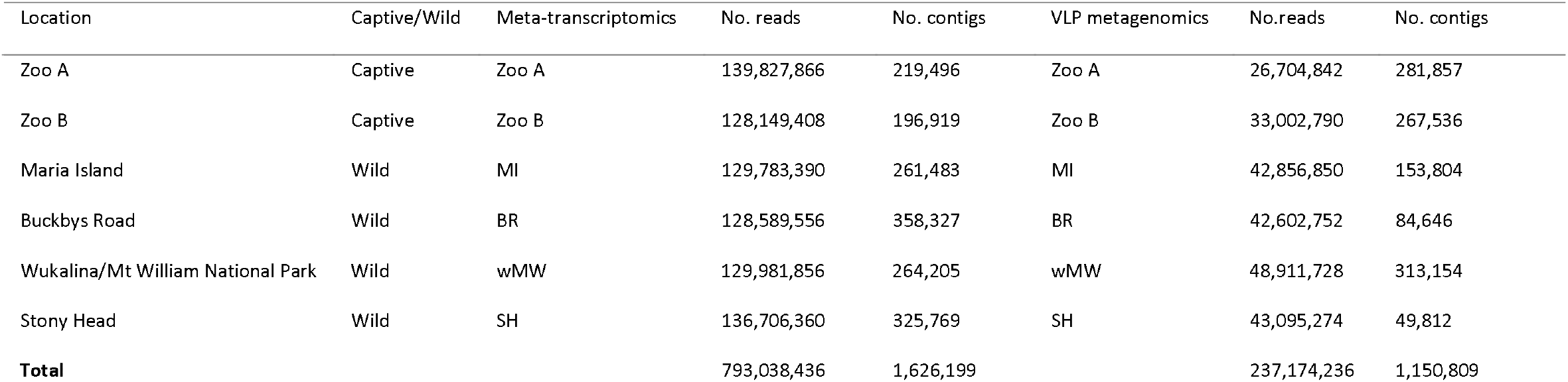
Library information of meta-transcriptomics and VLP metagenomics in the present study

|  |  |  |  |  |  |  |  |
| --- | --- | --- | --- | --- | --- | --- | --- |
| Zoo A | Captive | Zoo A | 139,827,866 | 219,496 | Zoo A | 26,704,842 | 281,857 |
| Zoo B | Captive | Zoo B | 128,149,408 | 196,919 | Zoo B | 33,002,790 | 267,536 |
| Maria Island | Wild | MI | 129,783,390 | 261,483 | MI | 42,856,850 | 153,804 |
| Buckbys Road | Wild | BR | 128,589,556 | 358,327 | BR | 42,602,752 | 84,646 |
| Wukalina/Mt William National Park | Wild | wMW | 129,981,856 | 264,205 | wMW | 48,911,728 | 313,154 |
| Stony Head | Wild | SH | 136,706,360 | 325,769 | SH | 43,095,274 | 49,812 |
| **Total** |  |  | **793,038,436** | **1,626,199** |  | **237,174,236** | **1,150,809** |

**Figure 2.**
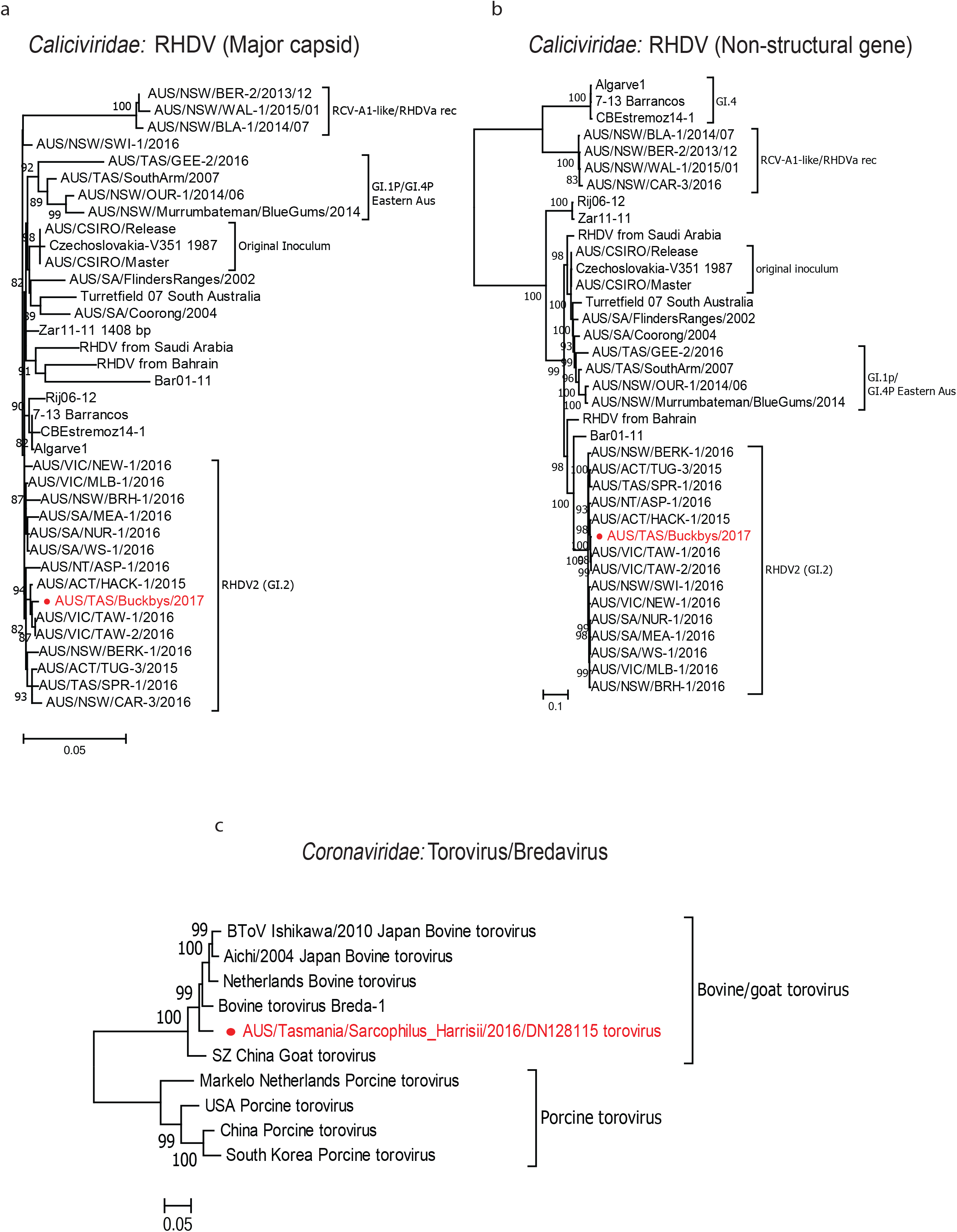
Overview of the Tasmanian devil faecal virome characterized by meta-transcriptomics and VLP metagenomics. Sequencing library/sampling site are represented at the bottom of the bar charts by SH for Stony Head, BR for Buckbys Road, wMW for wukalina/Mt William National Park, MI for Maria Island and Zoo A and B for the two captive populations. **a)** Proportions of all sequence reads in meta-transcriptomics (top) and VLP metagenomics (bottom) showing the proportions of reads belong to bacteria, eukaryote, archaea, viruses, host and unidentified. **b)** Proportions of RNA and DNA viruses detected by meta-transcriptomics (top) and VLP metagenomics (bottom). **c)** Virome composition and the proportions of viral groups in meta-transcriptomics (top) and VLP metagenomics (bottom). **d)** Estimated counts as calculated by RSEM (all six sequencing libraries from both VLP metagenomics and meta-transcriptomics combined and log transformed) to a selection of viruses, showing the differences in viruses detected by VLP metagenomics (blue) and meta-transcriptomics (orange).

VLP metagenomics resulted in 26 – 49 million reads per pool (237,174,236 reads in total), which were assembled *de novo* into 49,813 – 313,354 contigs (Table 1). In comparison to meta-transcriptomics, the VLP metagenomics protocol resulted in smaller proportions of non-viral nucleic acids (host and bacterial) and hence enriched viral nucleic acids (Figure 2a). The proportion of viral reads using VLP metagenomics varied from 14.69 – 60.02%, while the proportion of reads mapped to non-viral components were 17 – 49.46% for Bacteria, 0.17–2.42% for Eukarya, 1.29 –18.67% for host and less than 0.01% for Archaea (Figure 2a). For both approaches, a substantial proportion of reads had no significant similarity to other sequences in the databases on GenBank (10.22 – 29.20%).

Despite detecting a smaller proportion of virus-related sequences with meta-transcriptomics than with VLP metagenomics, viruses detected with the meta-transcriptomics approach fell into a wide range of viral groups, of which 49.87 – 97.51% had the closest hits to RNA viruses and 2.49 – 50.13% to DNA viruses. Conversely, for VLP metagenomics, over 95.54% of the virus-related sequences had closest hits to DNA viruses, and less than 5% were identified as RNA viruses (Figure 2b). Meta-transcriptomics revealed high levels of viral diversity across all libraries, with the most abundant viral groups detected being the *Caudovirales, Luteo-Sobemo, Narna-Levi, Partiti-Picobirna, Picorna-Calici* and *Tombus-Noda* (Figure 2c). Conversely, VLP metagenomics revealed relatively lower viral diversity across the same libraries, with *Caudovirales* dominating significant proportions of the viral reads (69.89-99.49%). Other viral groups identified by VLP metagenomics, at much lower abundances, include the *Microviridae, Circoviridae, Genomoviridae, Parvoviridae, Herpesviridae, Polyomaviridae and Papillomaviridae* (Figure 2c). Overall, VLP metagenomics and meta-transcriptomics differed in the viruses detected as well as the expected counts (transcript abundance) as measured by the RSEM analysis (Figure 2d).

The percentage of vertebrate viruses detected by meta-transcriptomics ranged between 0 –9.41% of the total viral reads, while large proportions of viral reads belonged to either non-vertebrate eukaryotic viruses (45.08 – 97.51%), which included plant viruses, insect viruses and mycoviruses, or bacteriophage (2.48 – 48.91%) from the families *Siphoviridae, Podoviridae, Myoviridae and Microviridae*. In the VLP metagenomics data set, the percentage of reads associated with vertebrate viruses was also small, ranging between 0.04 – 0.84%, while the percentages of bacteriophage and other eukaryotic viruses ranged between 79.17 – 99.91% and 0.04 – 19.99%, respectively. Detailed information on all vertebrate viruses identified is presented in Additional file 1: Table S1.

## Detection of viruses previously identified in other mammalian hosts (Rabbit haemorrhagic disease virus and Torovirus)

Rabbit haemorrhagic disease virus (RHDV) is a calicivirus in the genus *Lagovirus* [34]. All lagoviruses have a single-stranded, positive-sense RNA genome of approximately 7.5 kb and share a similar genome structure comprising two open reading frames (ORFs). Using meta-transcriptomics, we detected genomes with high nucleotide and amino acid similarity (>98 %) to RHDV in one of the wild devil meta-transcriptomic libraries (BR), with genome coverage of 98.1%. Phylogenetic analysis based on the nucleotide sequences of the major capsid and non-structural protein genes revealed that the RHDV detected here clustered with RHDV strain GI.2 (also called RHDV2) (Figure 3a and b), which was first detected in Australia in May 2015 and has since became the dominant circulating strain nationwide. RHDV-specific RT-PCR and sequencing confirmed the presence of RHDV2 in 4 of the 9 devil faecal samples from the BR meta-transcriptomics pool. In addition, no rabbit associated genes were detected during the initial sequence analysis. Additional PCR targeting a short fragment of rabbit mtDNA (<300bp) also did not detect any rabbit DNA in the original faecal samples from BR. Further RT-PCR and sequencing performed on the faecal RNA extractions from the remaining pools confirmed the presence of RHDV in 1 of 10 devils from wMW, 2 of 9 devils from Zoo A and 1 of 9 from Zoo B. One of the 4 additional RHDV positive samples, from Zoo A contained detectable levels of rabbit mtDNA as confirmed by PCR.

Torovirus is a genus of viruses in the viral family *Coronaviridae* with a linear, positive-sense RNA genome of about 28 – 28.5kb [38]. We identified the complete viral genome (28,463 bp) of a novel torovirus most closely related to Bovine torovirus (Bredavirus) in one of the meta-transcriptomic libraries (wMW). BLAST search revealed that the torovirus detected here shared 96% nucleotide similarity and 97% amino acid similarity with Bovine torovirus, a respiratory and enteric pathogen of cattle that causes gastroenteritis and severe diarrhoea, particularly in young calves. We determined the full genome structure of the novel torovirus strain, which included ORF 1a and ORF 1b encoding the two polyproteins (pp1a and pp1ab), ORF 2 encoding for the spike protein (S), ORF 3 encoding the membrane protein (M), ORF 4 encoding the hemagglutinin-esterase protein (HE), and ORF 5 encoding the nucleocapsid protein (N) [39].

**Figure 3.**
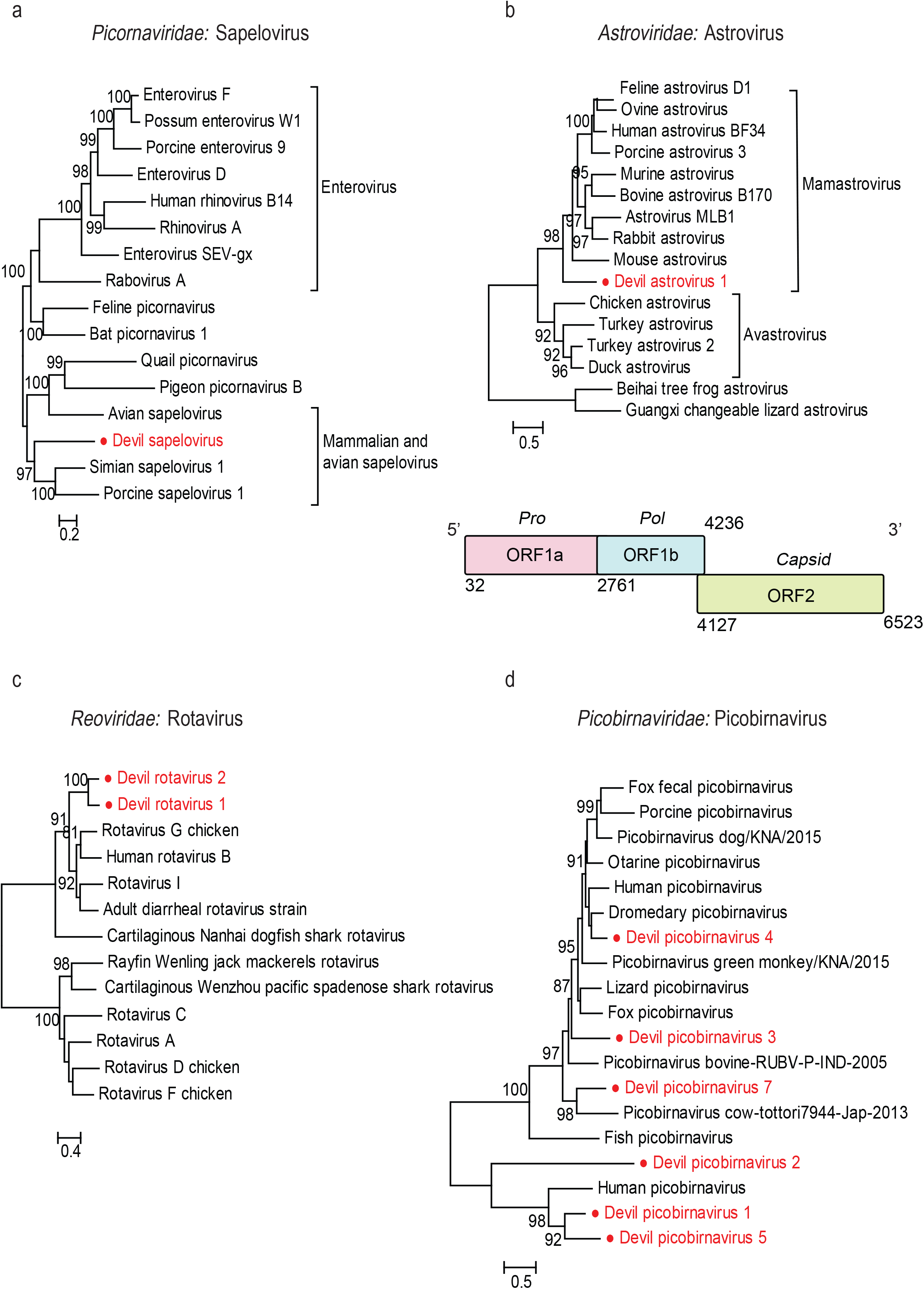
Maximum likelihood phylogenetic trees of viruses detected that were previously identified in other mammalian hosts. **a)** phylogenetic analysis of representative RHDV strains based on the 1414 nt nucleotide sequence of major capsid protein. **b)** phylogenetic analysis of representative RHDV strains based on the 5890 nt nucleotide sequence of the non-structural protein. **c)** phylogenetic analysis based on the 4762 aa spike protein of the novel torovirus strain detected here with other previously identified toroviruses. All trees are mid-point rooted and scaled to either the number of amino acid substitution or nucleotide substitutions per site based on the nature of alignment. The viral sequences detected in this study are shown in red in each tree.

Based on the phylogenetic analysis of the spike protein amino acid (aa) sequence (4762 aa), clustering of the novel torovirus strain with other toroviruses isolated from cows in the USA, Japan and Europe indicated a bovine origin, although this will need to be confirmed with wider sampling (Figure 3c).

## Detection and characterisation of novel marsupial-associated viruses (*Picornaviridae, Astroviridae, Reoviridae, Picobirnaviridae, Parvoviridae, Papillomaviridae, Herpesviridae, Polyomaviridae* and *Circoviridae*)

The order *Picornavirales* includes a diverse group of vertebrate-infecting RNA viruses, including enteroviruses and rhinoviruses, that can cause a wide range of diseases, such as poliomyelitis, hand, foot and mouth disease, encephalitis, respiratory tract infections and the common cold [40]. Members from the order are small, non-enveloped, positive-sense, single-stranded RNA viruses about 7-8.8 kb in size [41]. There are currently five recognized families: *Picornaviridae, Secoviridae, Iflaviridae, Marnaviridae* and *Dicistroviridae* [39]. We identified a novel member of the *Picornaviridae* in one of the meta-transcriptomic libraries (wMW) and obtained the complete genome of 8015 bp. According to the ICTV, members of a *Picornavirus* genus should share at least 40% amino acid sequence identity in the polyprotein region [39]. The encoded 2396 aa polyprotein exhibited 45.5% amino acid similarity to Simian sapelovirus, placing it in the genus *Sapelovirus*. We have provisionally named this newly identified virus Devil sapelovirus (DeSV). Phylogenetic analysis based on the amino acid sequence of the RdRp domains showed that DeSV formed a sister lineage to sapeloviruses identified from eutherian mammals (i.e. Porcine and Simian sapelovirus) (Figure 4a).

**Figure 4.**
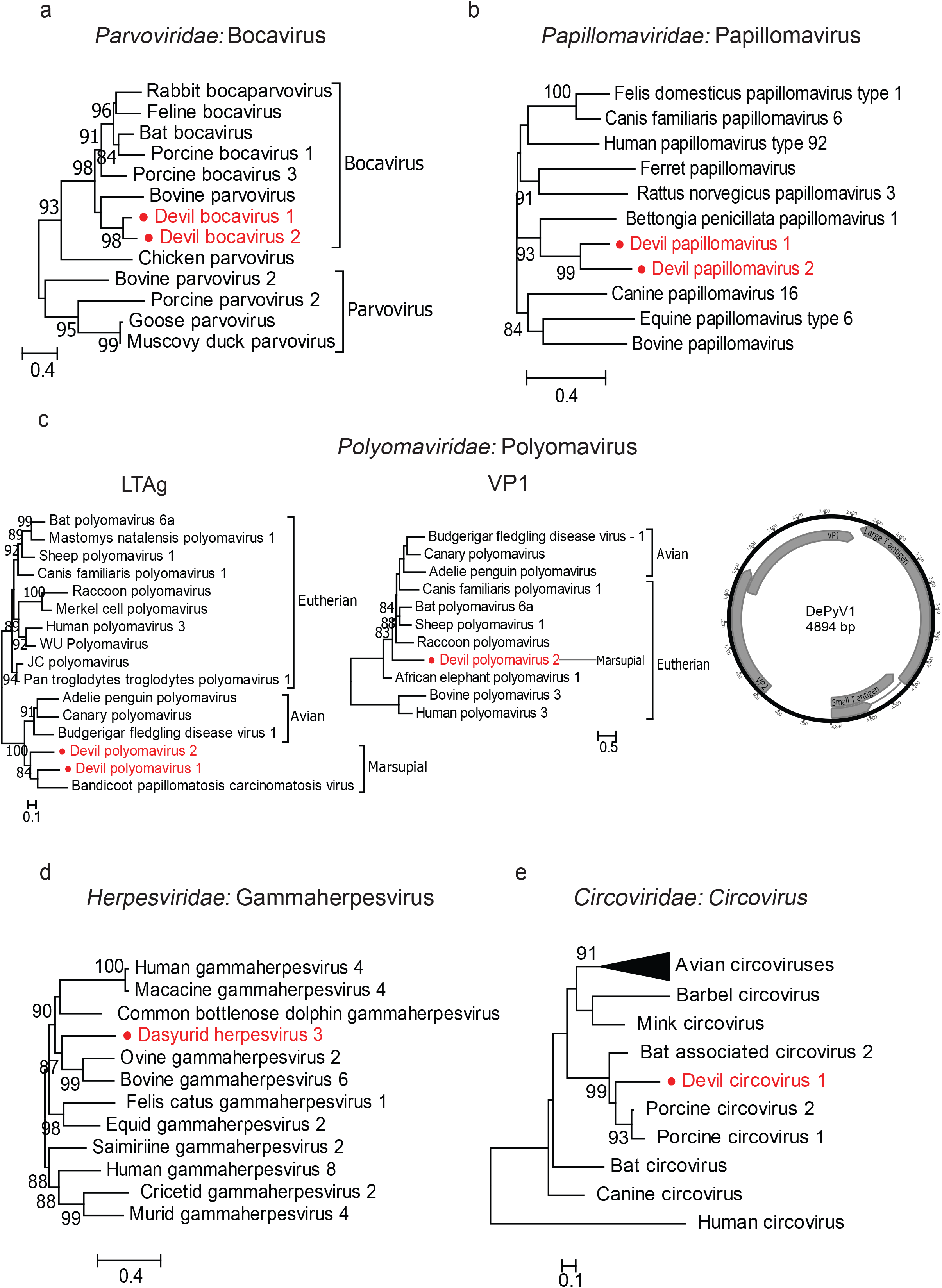
Phylogenetic analyses and genomic structures of the RNA viruses identified in the faeces of Tasmanian devils. All phylogenetic analyses were performed based on the amino acid sequence of the RdRp. **a)** Devil sapelovirus, **b)** Devil astrovirus, **c)** Devil rotavirus 1 and 2 and **d)** Devil picobirnavirus 1-6. For Devil astrovirus where the whole genome sequence was obtained, the genomic structure was also included and are shown below the corresponding phylogenetic tree. Predicted ORFs of these genomes are labelled with information of the potential protein or protein domain they encode. All trees are mid-point rooted and scaled to the number amino acid substitutions per site. The newly discovered viruses are shown in red in each phylogenetic tree.

Astroviruses are single-stranded positive-sense RNA viruses from the family *Astrovirida*e, that can cause diarrhoea and gastroenteritis in infected hosts [42, 43]. The family has a genome size of 6.4 to 7.3kb and has been identified in a broad range of vertebrate hosts including humans, other mammals, reptiles, amphibians, fish and birds [44]. We detected astrovirus-related sequences in 5 of the 6 meta-transcriptomic libraries (SH, wMW, BR, Zoo A and Zoo B). We identified one complete and one near-complete genome sequence with 81.4% pairwise nucleotide identity, denoting 2 separate strains of a single astrovirus species, which we have tentatively named Devil astrovirus 1 (DeAstV1) [45]. The DeAstV1 identified here has a genome structure typical of other astroviruses, with 3 putative open reading frames (ORF1a, ORF1b and ORF2) each encoding for the protease, RdRp and capsid, respectively. In addition, we found a ribosomal frameshift motif (AAAAAAC) within the ORF1a/1b overlap region. Phylogenetic analysis based on the conserved RdRp domain showed that DeAstV1 formed a distinct cluster that is more closely related to astroviruses of mammalian hosts (Mamastroviruses) than those of avian hosts (Avastroviruses) (Figure 4b).

Rotaviruses of the family *Reoviridae* are non-enveloped, double-stranded RNA viruses that infect only vertebrates through faecal-oral transmission [46]. They are the most common cause of acute viral gastroenteritis and can be found in a wide range of vertebrate hosts, predominately terrestrial mammals and birds. We detected rotavirus sequences in 3 of 6 meta-transcriptomic libraries (SH, BR and wMW, all are wild sites). Among them, we were able to identify 2 distinct segments (3481 bp and 3479 bp in length) encoding rotavirus VP1 (i.e. RdRp) from the two meta-transcriptomics libraries (BR and wMW, respectively). The two RdRp sequences shared >90 % nucleotide identity with each other, indicative of two different strains belonging to the same species. In addition, a contig encoding a partial rotavirus VP1 of 282 aa sharing 51 % sequence similarity with rotavirus H was also detected in meta-transcriptomics library wMW. We provisionally named the two viruses Devil rotavirus 1 and Devil rotavirus 2 (DeRoV1 and -2). The VP1 of DeRoV1 shared the highest amino acid similarity of ~51% with Rotavirus G, while DeRoV2 shared the highest amino acid similarity of 44% with Rotavirus H. Phylogenetic analysis with other previously characterised rotavirus species based on VP1 suggested that DeRoV1 and 2 form a distinct cluster that is most closely related to the cluster that contains Rotavirus G chicken, Human rotavirus B, Rotavirus I and Adult diarrheal rotavirus strain, which are associated with avian and mammalian hosts (Figure 4c).

Picobirnaviruses (PBV) are small, positive-stranded non-enveloped RNA viruses with abroad host ranges found in the faeces of many mammal, bird and reptile species [47]. We detected picobirnavirus sequences that encoded complete and partial viral RdRp (330 – 557 aa) in 4 of the 6 meta-transcriptomics libraries, all of which comprised of faecal samples collected from wild devils (BR, wMW, SH and MI). However, picobirnavirus sequences detected in library MI were too short to be phylogenetically informative and were discarded in the phylogenetic analysis. We provisionally named the novel picobirnaviruses detected here Devil picobirnavirus 1, 2, 3, 4, 5, and 6 (DePBV1 -6) with two separate strains occurring in DePBV1 and DePBV5. Phylogenetic analysis based on the RdRp domain of these viruses with other representative members from the family showed that the 6 novel Devil picobirnaviruses are highly diverse and widely distributed across the phylogenetic tree (Figure 4d).

*Parvoviridae* are a family of small, non-enveloped, ssDNA viruses [39]. Members of *Parvoviridae* are associated with a number of diseases, including erythema infectiosum in humans, hepatic and enteric diseases across a broad host range [48]. We identified two new members of the vertebrate-associated sub-family *Parvovirinae* in the faeces of Tasmanian devils from two of the VLP metagenomics libraries (MI and BR). We recovered partial and near-complete protein sequences sharing ~50% identity to California sea lion bocavirus and Porcine bocavirus, respectively. Two bocavirus-related sequences detected here shared >97% amino acid sequence similarity, denoting two separate strains of the same species, which we have provisionally named Devil bocavirus 1 (DeBoV1). A third bocavirus-related sequence was also identified in VLP metagenomics library BR, sharing 71.13% amino acid similarity with DeBoV1. We provisionally named this second virus Devil bocavirus 2 (DeBoV2). Phylogenetic analysis of DeBoV1 and 2 in the context of other representative viruses from the *Parvoviridae* family further confirmed the clustering of DeBoVs within the diversity of mammalian bocaviruses, although the branching order involved DeBoVs and other members of bocavirus remain unresolved with current topology (Figure 5a).

**Figure 5.**
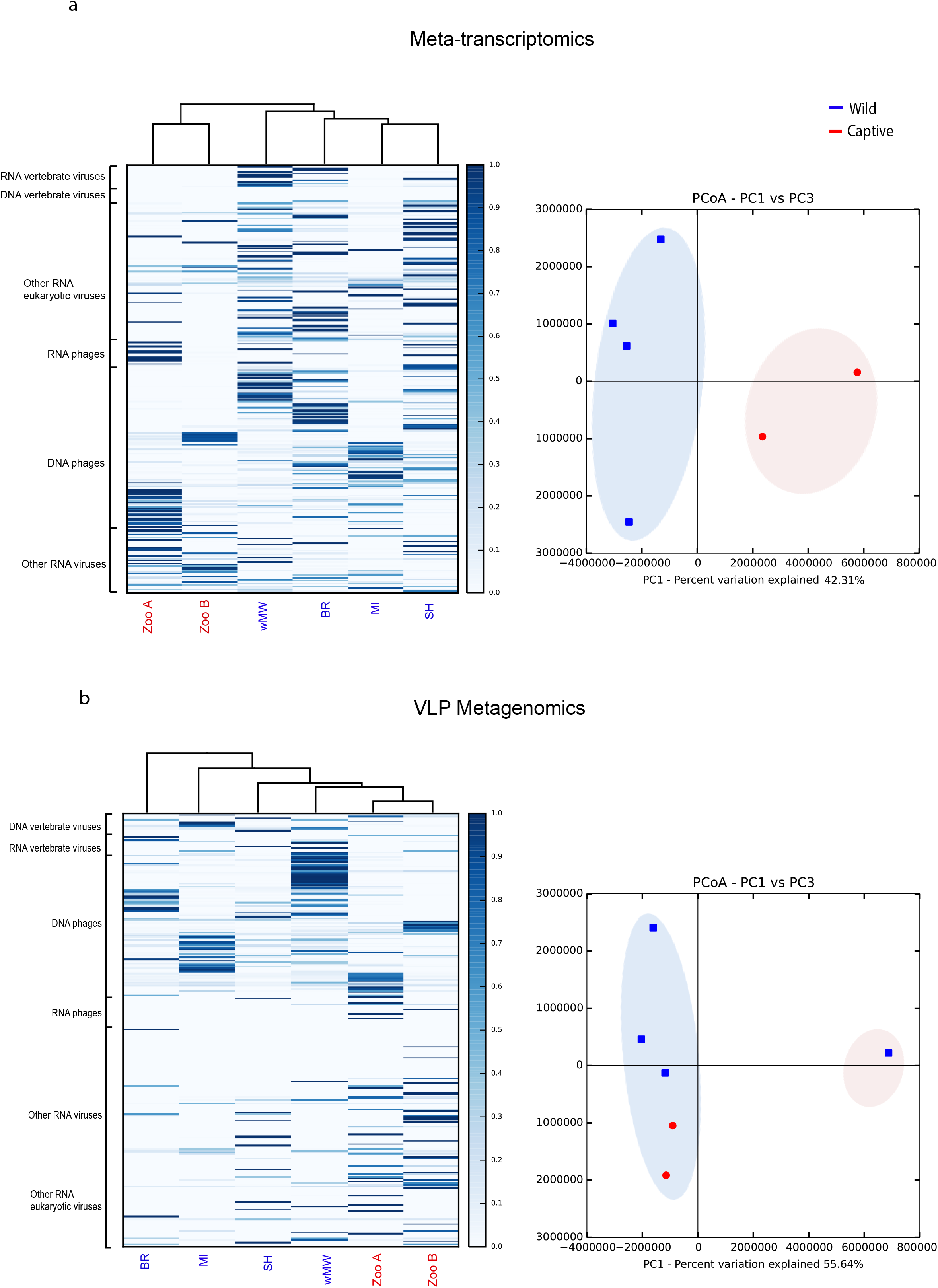
Phylogenetic analyses and genomic structures of the DNA viruses identified in the faeces of Tasmanian devils. **a)** Devil bocavirus 1 and 2 based on the amino acid sequences of the NS1 protein. **b)** Devil papillomavirus 1 and 2 based on the amino acids of the E1 protein. **c)** Devil polyomavirus 1 and 2 based on the amino acid sequences of the LTAg and VP1 proteins. For devil polyomavirus 1, the whole genome sequence was obtained and the genomic structure is shown next to the VP1 tree. Predicted ORFs of these genomes are labelled with information of the potential protein or protein domain they encode. **d)** Dasyurid herpesvirus 3 based on the amino acid sequence of the DNA polymerase. **e)** Devil circovirus 1 based on the amino acid sequence of the replicase protein. All trees are mid-point rooted and scaled to the number of amino acid substitutions per site. The newly discovered viruses are shown in red.

Papillomaviruses from the family *Papillomaviridae* are small, nonenveloped, icosahedral viruses with a circular double-stranded DNA genome of about 8 kbp in size [39], and are associated with the development of benign and malignant tumours [49, 50]. We identified fragmented genome of two novel species of papillomavirus from a single VLP metagenomics library (MI), among which we were able to retrieve two longer fragments of 1225 and 1335 bp in length, respectively, both of which encode partial E1 protein, an ATP-dependent DNA helicase required for viral replication [51]. Since the two fragments shared approximately 64% similarity to each other, it suggests the presence of two distinct papillomavirus species, which we have tentatively designated as Devil papillomavirus 1 and 2 (DePV1 and -2). Phylogenetic analysis based on the E1 protein showed that DePV1 and 2 form a distinct cluster with *Bettongia penicillata* papillomavirus type 1 (BpPV1) isolated from the woylie (Figure 5b). While the marsupial papillomaviruses viruses are clustered together in the phylogenetic tree, their relationship with viruses identified from eutherian mammals remain unresolved.

Polyomaviruses from the family *Polyomaviridae* are small, nonenveloped, circular DNA viruses about 5 kb in size that have been identified in a wide variety of mammalian and avian hosts [39, 52]. We detected two novel polymaviruses from the faeces of Tasmanian devils in 3 VLP metagenomics libraries (MI, wMW and Zoo B), which we have tentatively designated Devil polyomavirus 1 and 2 (DePyv1 and -2), respectively. We recovered the complete circular genome of 4894 bp for DePyv1 (Figure 5c) and partial gene sequence (2251 bp) of the large T antigen (LTAg) protein for DePyv2. Phylogenetic analyses revealed strikingly different evolutionary histories for the structural and non-structural parts of the genome (Figure 5c), indicative of recombination [53]. In the LTAg phylogeny, DePyV1 and -2 formed a distinct lineage with another marsupial virus Bandicoot papillomatosis carcinomatosis virus type 2 (BPCV-2), which in turn is clustered with polyomaviruses of avian hosts (Figure 5c). In contrast, in the VP1 phylogeny, the marsupial viruses showed no close relationship with the avian viruses. Interestingly, the Bandicoot papillomatosis carcinomatosis viruses (BPCV1 and -2) only showed close relationship to DePyV1 at the LTAg region but not the VP1 region. Indeed, both BPCV1 and -2 are hybrid viruses between the family *Papillomaviridae* and *Polyomaviridae* [54, 55]. While its non-structural genes (i.e. large T and small T antigen genes) are related to polyomavirus, its structural genes (i.e. L1 and L2 protein genes) are related to papillomavirus [54].

*Herpesviridae* are a large family of DNA viruses with large linear, dsDNA genome of about 120-240 kb, and that comprise 3 subfamilies; *Alphaherpesvirinae*, *Betaherpesvirinae* and *Gammaherpesvirinae* [39]. In one of the captive VLP metagenomics libraries (Zoo A), we identified more than 70 contigs matching different regions of a herpesvirus genome, which totaled 62,821 bp in length and included partial gene sequences of the DNA polymerase (575 aa), major DNA binding protein (465 aa), helicase (396 aa), glycoprotein M (378 aa) and H (427 aa), and major capsid protein (358 aa), amongst others. We tentatively named the novel herpesvirus Dasyurid herpesvirus 3 (DaHV-3). Phylogenetic analysis based on these non-structural and structural proteins suggested that DaHV-3 clustered with other gammaherpesviruses from the *Gammaherpesvirinae* subfamily (Additional file 5: Figure S1 and Additional file 6: Figure S2). Further analysis of the *Gammaherpesvirinae* phylogenetic structure based on the DNA polymerase showed that DaHV-3 forms a distinct lineage that is most closely related to Bovine gammaherpesvirus 6 and Common bottlenose dolphin gammaherpesvirus 1 strain Sarasota (Figure 5d). The previously characterised Dasyurid herpesvirus 2 (DaHV-2) isolated from Tasmanian devils [12] could not be included in the phylogenetic analysis because there are no available sequences from the same genomic regions. A BLASTx search of the DNA polymerase showed that DaHV-3 exhibited the greatest amino acid similarity (93.3%) with the previously characterised Macropodid herpesvirus 3(MaHV-3) [12], whose DNA polymerase amino acid sequence was also too short (<50% of the other representative herpesviruses) to be included in the phylogenetic analysis.

Circoviruses from the family *Circoviridae* have a circular ssDNA genome which range from 1.7 to 2.3 kb in size [56]. Here, we identified circovirus-related sequences and recovered the partial replicase gene sequences (899 bp) in one of the wild devil metagenomes (SH) and tentatively named it Devil circovirus (DeCV). Phylogenetic analysis based on the Rep proteins of the novel DeCV and representative strains of circoviruses and cycloviruses suggested that DeCV is clustered with circoviruses previously isolated from bats and pigs, sharing the highest sequence identity (62%) with bat circovirus (AIF76281.1) (Figure 5e).

## Other viruses: plant and insect viruses, and bacteriophage

In both the VLP metagenomics and meta-transcriptomics analyses, large proportions of viral sequence reads could be attributed to viruses that infect plants, insects and bacteria. Indeed, bacteriophage sequences from the order *Caudovirale*s were detected in all libraries and made up over 90% of all viral VLP metagenomic reads and up to 48.91% of the meta-transcriptomic reads. Sequences related to newly identified arthropod viruses, such as Wuhan fly virus and Wuhan mosquito virus were also detected. Most of the insect viruses detected belong to RNA virus groups the *Bunyavirales*, the *Mononegavirales*, and the *Chuviridae*, as well as DNA virus subfamily *Densovirinae* from the *Parvoviridae* family. The detection of these arthropod virus related sequences indicates the possible ingestion of arthropods and/or the environmental contamination of faeces. Despite being a carnivorous species, sequences related to various plant and fungal viruses were also observed in all libraries, including various sobemoviruses, tombusviruses and mitoviruses. Herbivorous species such as wallabies often make up a large proportion of devil’s diet, and the presence of plant viruses in these prey items might be easily detected in the faeces of devil through deep sequencing, especially if the intestines of the prey were consumed [1]. Nonetheless, devils have also been observed to consume vegetation, and the abundance of highly diverse eukaryotic and vertebrate viruses that are thought to be associated with diverse host taxa are reflective of devils’ overall generalist diet [1].

## Virome ecology: comparison between devil populations

Within-library viral diversity (alpha diversity) as characterised by our meta-transcriptomics approach was significantly different between captive and wild populations (*p* < 0.05). In general, captive populations had lower diversity in their faecal viromes compared to wild populations, as measured using the chao1 metric. However, when considering bacteriophage alone, we did not find any significant associations between devil populations (captive or wild) in either diversity or abundance.

The meta-transcriptomics analysis of the virome of Maria Island (MI) devils displayed a similar level of overall viral diversity to the two captive populations (Zoo A and Zoo B), and was lower than that found in the three other wild populations (wMW, BR and SH) (Figure 6a). Conversely, alpha diversity as characterized from VLP metagenomics data did not differ significantly between libraries (Figure 6b). Cluster analysis indicated that in the meta-transcriptomics data, wild and captive devils fell into two distinct clusters, while in VLP metagenomics data, BR formed its own cluster and the remaining populations formed a second cluster (Figure 6a and b).

**Figure 6.**
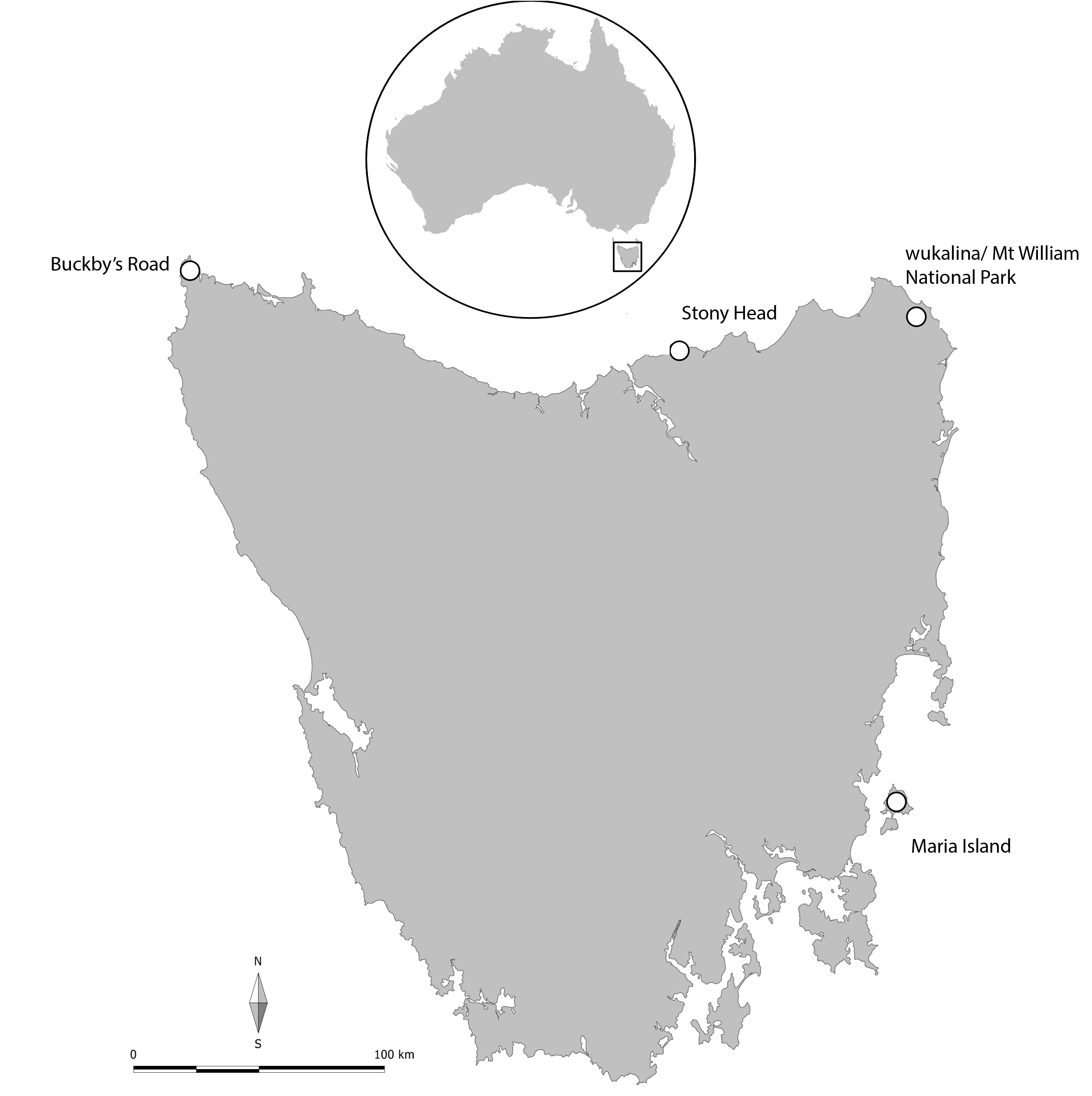
Heatmaps (left) showing the hierarchical clustering and percentage of reads from each library mapping to various viral groups, and principal coordinate analysis (PCoA) plots (right) showing the similarity relations among libraries based on Euclidean distances as seen in **a)** meta-transcriptomics and **b)** VLP metagenomics.

## Discussion

We provide the first comprehensive characterisation of the faecal virome of an endangered marsupial species, the Tasmanian devil. Using both VLP metagenomics and meta-transcriptomics, we identified a huge diversity of viruses in the faeces of devils, including vertebrate viruses, bacteriophage and other eukaryotic viruses. Novel viruses identified as potentially marsupial-associated – including, sapelovirus, rotaviruses, picobirnaviruses, astrovirus, bocaviruses, papillomaviruses, herpesvirus and polyomaviruses – were detected and have greatly expanded our current knowledge of viruses that are found within this unique group of mammals.

Our comparison of VLP metagenomics and meta-transcriptomics approaches revealed marked differences in terms of the viruses detected. In general, VLP metagenomics mainly detected DNA viruses, while meta-transcriptomics detected both DNA and RNA viruses, although the DNA viruses detected with meta-transcriptomics were limited to those with relatively high abundance (Fig 2d). A high abundance level is often indicative of an active viral infection, during which DNA viruses are transcribed into RNA intermediates that can be readily detected by RNA sequencing [57]. Conversely, RNA viruses identified in meta-transcriptomic were rarely detected in VLP metagenomics, even if they were highly abundant based on the RSEM estimated counts.

The use of VLP enrichment and sequence-independent amplification in VLP metagenomics increased the number of viral reads in each library compared to the meta-transcriptomic approach. However, the viral compositions of all six VLP metagenomes were highly skewed towards DNA viruses, particularly DNA bacteriophage from the order *Caudovirales*. This is consistent with a previous study comparing various enrichment methods, in which bacteriophage accounted for >80% of all reads in all of the enrichment methods tested, but <5% when no enrichment steps were incorporated [58]. Despite being able to substantially increase the total number of viral reads in the metagenomes, sequence-independent amplification is bias prone, resulting in fewer viruses being detected and lower genome coverage due to preferential amplification of certain sequences [59-61]. However, the over-representation of bacteriophage found here could also be attributed to the fact that they make up the bulk of the virobiota in the gut, which is dominated by bacteria [62, 63]. Regardless of its known bias [58, 59], VLP metagenomics still holds merit for use in virome characterisation due to its ability to identify low abundance DNA viruses, which is especially relevant for dormant or non-active viruses.

Compared to VLP metagenomics, meta-transcriptomics is non-viral specific, requires less sample processing, and reveals the entire transcriptome within a sample [20, 64]. Omitting the need for VLP enrichment and additional sample processing, the likelihood of biased detection is plausibly reduced in meta-transcriptomics. In this study, the proportions of viral reads sequenced by meta-transcriptomics were less than 2% per library, but the numbers of viral groups detected were significantly higher than those detected in VLP metagenomics, which included both RNA and DNA viruses.

Importantly, then, our results show that the taxonomic compositions of viral communities as revealed by VLP metagenomics and meta-transcriptomics were not interchangeable and neither of the approaches was able to detect all viruses present. However, these two approaches were complementary to one-another, and an integrated approach using both VLP metagenomics and meta-transcriptomics will prove to be a powerful tool for obtaining a complete overview of both the taxonomic and functional profiles of viral communities in a sample.

Ecological analysis of virome composition and diversity revealed significant differences between captive and wild devil populations, especially when characterised using meta-transcriptomics. The two captive populations displayed lower levels of viral diversity compared to the four wild populations. This loss of diversity within the captive populations is consistent with trends previously observed in the gut bacteriome, where captive devils also exhibited lower bacterial diversity compared to wild devils [65]. The microbiome is dynamic and sensitive to changes in the environment. Hence, the extreme lifestyle and diet changes that often occur in captivity are likely to affect the viral communities of devils. Interestingly, Maria Island devils had viromes that are more similar to the two captive populations. Maria Island is a 115 km^2^ island off the east coast of Tasmania (Fig 1) and is home to approximately 100 free ranging devils. There were no Tasmanian devils living on Maria Island until 27 devils were introduced to the island in 2012 and 2013 from several captive populations [66]. Non-native vertebrate species, such as cats, black rats and house mice, are present on the island, whilst other species such as rabbits and livestock are absent, although all these species are common in the wild on mainland Tasmania. Two factors may have potentially contributed to the lower viral diversity observed on Maria Island. First, due to its isolation from mainland Tasmania, animal movements or immigrations are limited to only marine or bird species. This restriction to animal movements (particularly vertebrate species) between the mainland and Maria Island may in turn limit the introduction of viruses to the island. Second, captive-born devils are likely to have a “captive-type” virome, that is, lower viral diversity compared to wild devils. Indeed, some of the devils included in this study are captive-born animals recently translocated to the island. As samples were pooled prior to sequencing, it was not possible to distinguish between viromes of captive-born and wild-born devils. To understand whether captive devils can reacquire a “wild-type” virome following translocations, future studies should focus on comparing captive born translocated devils with the incumbent devils in the wild.

Overall, we detected sequences related to 20 vertebrate viruses, including 18 novel marsupial related viruses and two known mammalian viruses (RHDV2 and torovirus). While some of the viruses identified in this study come from families that include important pathogens, their pathogenic potential in devils is unclear. It is also important to note that some of these viruses may in fact be dietary viruses or occur naturally as part of the normal gut flora and therefore do not necessarily lead to disease. In the case of RHDV, we showed that RHDV2 was present in both captive and wild devils. In some areas of Tasmania, rabbits are common and may be a dietary component of wild devils. They are also regularly fed to devils in captivity, along with other meat sources such as wallaby and chicken. Feeding records provided by the two zoos in this study confirmed the feeding of rabbits to some of the sampled devils (zoo A) around the time of faecal collection. While we did not detect rabbit genes by meta-transcriptomics, VLP metagenomics or PCR in any of the wild devil samples that tested positive for RHDV, we did detect rabbit mtDNA in the stools of one captive devil from zoo A that tested positive for RHDV by PCR. Thus, more specific diagnostic techniques such as targeted PCR of blood or internal tissues (i.e. liver), *in situ* hybridization and serological assays are required to determine whether these viruses can actively replicate in devils and cause disease or are simply gut contaminants. Nevertheless, exposure to host-adapted viruses could pose significant health threats, especially for devils that are immunocompromised due to DFTD, old age or other concurrent diseases [67, 68]. Furthermore, even commensal or latent viral infections can be exacerbated or reactivated in immunocompromised hosts [69, 70]. For example, in the giant panda, while a number of viruses (e.g. papillomaviruses, picornaviruses and anelloviruses) were detected in both sick and healthy animals, the virus titres were much higher in the diseased individuals, indicating a compromised balance between host immune response and virus replication [71].

Tasmanian devils have low genetic diversity both across their genomes and at functionally important loci such as the major histocompatibility complex (MHC) [72-74]. This lack of diversity renders them particularly vulnerable to changes in the environment, including the emergence of new pathogens [75]. For instance, populations of Italian agile frogs with lower microsatellite diversity have been shown to be more susceptible to an emerging strain of Ranavirus (FV3-frog virus 3) and experience higher mortality rates than populations with higher diversity [76]. Similarly in cheetahs, a coronavirus-associated feline infectious peritonitis outbreak causing mass mortality in a captive breeding colony was linked to the species’ extreme genetic monomorphism, particularly at the MHC [77]. While none of the devils included in this study exhibited overt signs of disease at the time of sample collection, the viruses described here will provide a fundamental baseline of the normal Tasmanian devil faecal virome, which can be used as a reference for comparing healthy and diseased animals.

Phylogenetic analyses of the newly identified viruses, including divergent members of their respective viral families, provided us with insights into the evolutionary history of marsupial-associated viruses relative to viruses of eutherian mammals and other host taxa. Generally, a long-term relationship between viruses and hosts are expected for mammalian viruses [25]. Strong evidence for this lies in the observation that devil viruses are usually clustered with other marsupial viruses, as a marsupial-specific lineage that is distinct from the eutherian viruses, as observed in papillomavirus [78, 79], polyomavirus, and herpesvirus (Fig. 5). Furthermore, in several cases the branching order of viruses broadly reflects that of their hosts such that a general co-divergence can be inferred. For instance, in the phylogenies of *Picornaviridae* and *Astroviridae*, the devil (marsupial) viruses formed sister clade to eutherian viruses, which in turn are sister to avian viruses, consistent with the evolutionary history of the host. Although such a relationship is not observed in every virus phylogeny, a deep divergence between eutherian and marsupial viruses is typical of our dataset. This observation indicates that the timescale of virus evolution is very likely to reflect that of the hosts [25].

The gut virome is increasingly recognized as an integral component of the gut microbiome, and studies of the devil virome will continue to shine light on the biology and health of this iconic endangered species. For example, bacteriophage, which we have shown to dominate the devil’s faecal virome, can contribute to host health by maintaining the diversity and structure of the gut bacteriome through direct interactions with the bacterial communities. While the functions of bacteriophage on devil health remain to be determined, future studies will be able to exploit the extensive microbiomic data that is now available to answer important questions about the host-microbe relationship between devils and their microbiome [65].

## Conclusion

We provided the first comprehensive characterisation of the faecal virome of the Tasmanian devil, greatly expanding our knowledge of the diverse groups of vertebrate viruses observed in this endangered species. Similar to their bacterial microbiome, captive devils have significantly lower diversity in their faecal viromes compared to wild devils, likely reflective of their captive diets and lifestyle. Identification of vertebrate and marsupial-specific viruses in devils provides potential candidate viruses for future disease surveillance and routine screening as part of the broader conservation management of devils. However, future work will first need to focus on elucidating the pathogenic impact of these viruses on devil health. Finally, we showed that a combination of VLP metagenomics and meta-transcriptomics may be a more comprehensive virome characterisation approach that will encompass both DNA and RNA viruses.

## Abbreviations

SH: Stony Head
MI: Maria Island
BR: Buckbys Road
wMW: wukalina/Mt William National Park
VLP: Virus-like particles
RHDV: Rabbit haemorrhagic disease virus
DeSV: Devil sapelovirus
DeAstV1: DeRoV1 and -2 Devil rotavirus 1 and 2
DePBV1-6: Devil picobirnavirus 1-6
DeBoV1 and -2: Devil bocavirus 1 and 2
DePV1 and -2: Devil papillomavirus 1 and 2
DePyV1 and -2: Devil polyomavirus 1 and 2
DaHV-3: Dasyurid herpesvirus 3
DeCV: Devil circovirus

## Acknowledgements

Thank you to the field teams of the Save the Tasmanian Devil Program and the University of Sydney who collected faecal samples. Thanks also to N. Mooney for his assistance at the BR site, the two zoos (and their keeping staff) who provided access to faecal samples from the captive devils, J. Mahar for providing rabbit liver tissue and A.V. Lee for the map in Figure 1.

## Availability of data and material

Raw sequence reads generated in this study are available in NCBI SRA database under the BioProject ID PRJNA495667. Viral genomes described in detail here are available in the figshare repository https://doi.org/10.6084/m9.figshare.7185146.v3 and GenBank (submission currently under process).

## Authors’ contributions

RC, VB, MS, CH, CEG and KB designed the experiments. CH coordinated sample collection from all sites, with RC, CEG, CH participating in collection from the zoos, BR, SH (respectively). RC carried out the extractions and PCRs. RC, MS analysed the data. RC wrote the manuscript and all authors critically reviewed and edited the manuscript.

## Ethics approval and consent to participate

Wild Tasmanian devils were trapped by the Save the Tasmanian Devil Program staff under the Tasmanian Government Department of Primary Industries, Parks, Water and Environment Standard Operating Procedure “Trapping and handling wild Tasmanian devils”.

## Consent for publication

Not applicable

## Additional files

**Additional file 1: Table S1** List of vertebrate viruses identified by meta-transcriptomics and VLP metagenomics, including information on the sequence library, identification, classification, virus and strain name, sample type, genome length and percentage reads, Blastx hits to known viruses. (.xls)

**Additional file 2: Table S2.** Virus abundance table showing the proportion of reads mapped to viral contigs in each meta-transcriptomic library. (.xls)

**Additional file 3: Table S3.** Virus abundance table showing the proportion of reads mapped to viral contigs in each VLP metagenomics library. (.xls)

**Additional file 4: Table S4.** List of primer sets used for PCR confirmation of RHDV2. (.xls)

**Additional file 5: Figure S1.** Maximum likelihood phylogenetic tree of Dasyurid herpesvirus 3 in the context of representatives of the *Herpesviridae* family based on the amino acid sequences of glycoprotein M. The tree is mid-point rooted and scaled to the number of amino acid substitutions per site. The newly discovered Dasyurid herpesvirus 3 is shown in red. (.pdf)

**Additional file 6: Figure S2.** Maximum likelihood phylogenetic tree of Dasyurid herpesvirus 3 in the context of representatives of the *Herpesviridae* family based on the amino acid sequences of major capsid. The tree is mid-point rooted and scaled to the number of amino acid substitutions per site. The newly discovered DaHV-3 is shown in red. (.pdf)

